# When three traits make a line: Evolution of phenotypic plasticity and genetic assimilation through linear reaction norms in stochastic environments

**DOI:** 10.1101/034256

**Authors:** Torbjørn Ergon, Rolf Ergon

**Affiliations:** Centre for Ecological and Evolutionary Synthesis, Department of Biosciences, University of Oslo, P.O. Box 1066 Blindern, N-0316 Oslo, Norway; University College of Southeast Norway, Porsgrunn, Norway

**Keywords:** genetic redundancy, environmental change, value of information, uncertain/imperfect cues, life-history theory, optimality models, cue perception and reliability

## Abstract

Genetic assimilation results from selection on phenotypic plasticity, but quantitative genetics models of linear reaction norms considering intercept and slope as traits do not fully incorporate the process of genetic assimilation. We argue that intercept-slope reaction norm models are insufficient representations of genetic effects on linear reaction norms, and that considering reaction norm intercept as a trait is unfortunate because the definition of this trait relates to a specific environmental value (zero) and confounds genetic effects on reaction norm elevation with genetic effects on environmental perception. Instead we suggest a model with three traits representing genetic effects that respectively (i) are independent of the environment, (ii) alter the sensitivity of the phenotype to the environment, and (iii) determine how the organism perceives the environment. The model predicts that, given sufficient additive genetic variation in environmental perception, the environmental value at which reaction norms tend to cross will respond rapidly to selection after an abrupt environmental change, and eventually become equal to the new mean environment. This readjustment of the zone of canalization becomes completed without changes in genetic correlations, genetic drift or imposing any fitness costs on maintaining plasticity. The asymptotic evolutionary outcome of this three-trait linear reaction norm generally entails a lower degree of phenotypic plasticity than the two-trait model, and maximum expected fitness does not occur at the mean trait values in the population.

## Introduction

All natural populations evolve in environments that are to some degree variable. Biologists have long realized that the phenotypic expression of different genotypes may respond differently to the same environmental change, and that such phenotypic plasticity may be heritable (DeWitt & Scheiner, 2004; Pigliucci, 2005). Depending on the effect this phenotypic plasticity has on selection (fitness), evolution may thus bring about mechanisms that either buffer the phenotypic expression against environmental variation (i.e., environmental canalization) or modify the responses to some environmental influence in an adaptive manner (Nijhout, 2003). Phenotypic plasticity involves developmental, physiological and/or behavioral phenotypic responses to some component(s) of the environment (DeWitt & Scheiner, 2004; Pigliucci, 2005; Pigliucci *et al.*, 2006). These environmental components, often referred to as environmental ‘cues’(DeWitt & Scheiner, 2004), are often just correlated with, but not identical to, the environmental variables affecting fitness (e.g. McNamara *et al.*, 2011; Svennungsen *et al.*, 2011; Gienapp *et al.*, 2014). Hence, cues do not provide perfect information about the optimal phenotypic expression, and it is usually adaptive to respond more conservatively towards information-poor cues than more informative ones (Yoccoz *et al.*, 1993; Ergon, 2007; McNamara *et al.*, 2011). The phenotypic expression of a particular genotype as a function of environmental cues is called a reaction norm (Woltereck, 1909; Pigliucci, 2005). There has been considerable interest in evolutionary processes governing reaction norms as this is crucial for our understanding of how populations may respond to environmental change and introduction to novel environments (e.g. Lande, 2009; Reed *et al.*, 2010; McNamara *et al.*, 2011; Gienapp *et al.*, 2014).

Waddington (1953, 1961) originally used the term ‘genetic assimilation’ to describe experimental selection results where qualitative phenotypes (such as lack of cross-veins in *Drosophila* wings) that are initially only expressed in response to a particular environmental stimuli (such as heat shock during a particular stage of development) becomes constitutively produced (i.e., becomes expressed independently of the environmental stimuli) after continued selection. However, ‘genetic assimilation’ is also used to describe similar phenomena in evolution of the mean of quantitative phenotypes that may remain plastic at equilibrium in a stochastic environment after an environmental change (Pigliucci & Murren, 2003; Lande, 2009). In such cases, the new equilibrium phenotypes will not be independent of the environment unless the reaction norm slope is zero.

We here use the term ‘genetic assimilation’ essentially as in Pugliucci *et al.* (2006) and Lande (2009) to describe the evolutionary scenarios where, after an abrupt environmental change, there is an initial increase in phenotypic plasticity, after which mean plasticity is reduced and the zone of canalization (i.e., the environment range, or value, where phenotypic variance is at minimum; Dworkin, 2005; Lande, 2009) moves towards the current mean environment (see Fig. 2 in Pugliucci *et al.* (2006) and Fig. 1 in Pigliucci & Murren (2003)). While the exact definition of ‘genetic assimilation’ and proposed mechanisms are somewhat contentious (Scharloo, 1991; Pigliucci *et al.*, 2006), there is substantial evidence from both laboratory experiments and field studies that such processes do occur (Pigliucci & Murren, 2003; Braendle & Flatt, 2006; Pigliucci *et al.*, 2006). In our treatment, we regard the process of genetic assimilation as complete in a stationary environment when phenotypic variance is minimized in the mean environment (but both mean reaction norm slope and phenotypic variance in the mean environment may remain non-zero). Population level phenotypic variation in a fluctuating environment depends on both the degree of environmental canalization, or “buffering”, of individual plasticity (represented by the genotypic reaction norm slopes; Dworkin, 2005) and the variation among genotypes in the reaction norm elevation around the mean environment. In a population of linear reaction norms, phenotypic variance is always minimized in the environment where the correlation between reaction norm slope and the phenotypic expression is zero (i.e., where reaction norms “tend to cross”; Lande (2009)).

**Figure 1.**
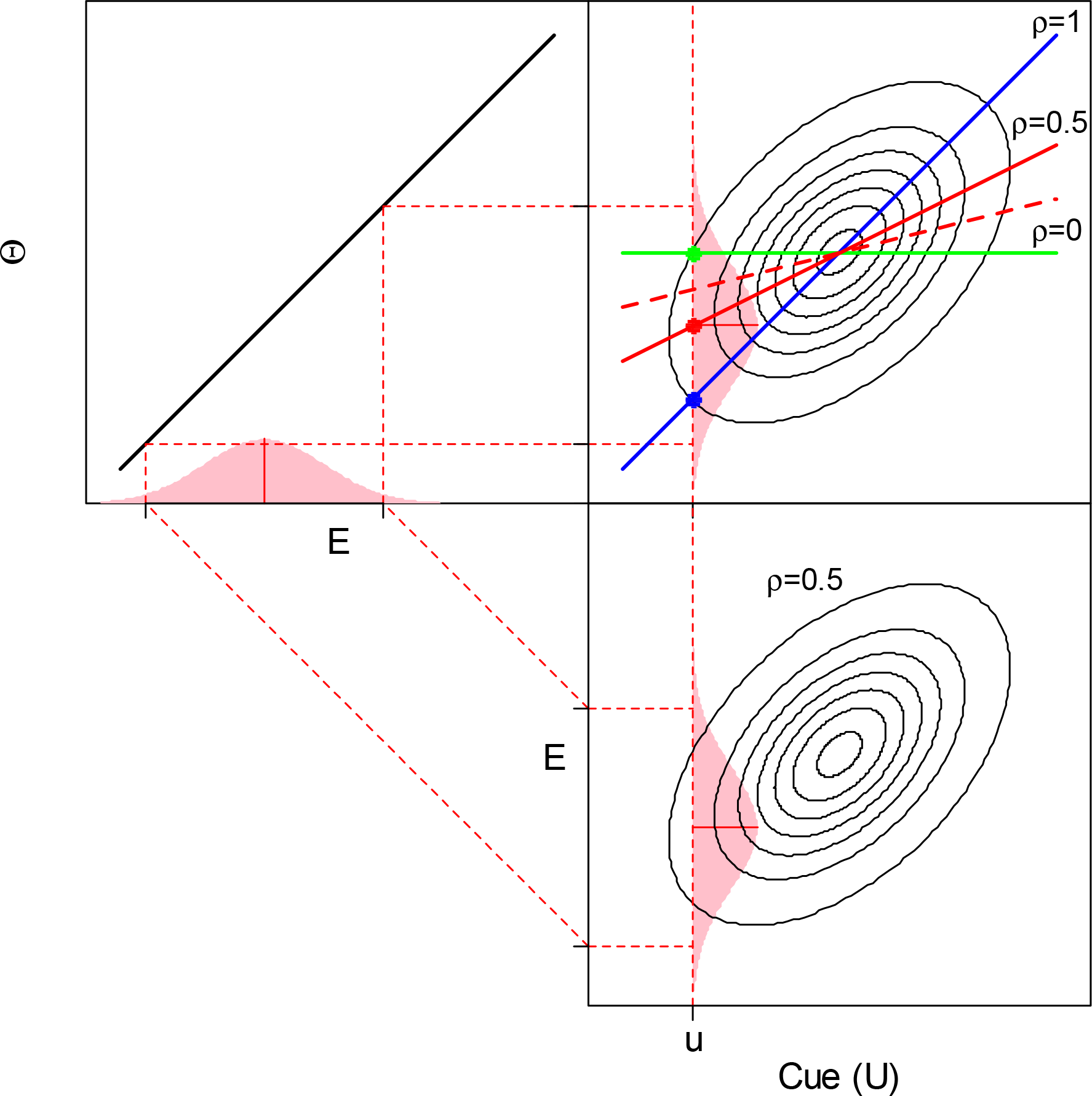
Conceptual overview of optimal linear reaction norms in stochastic environments. The environmental component *U* (cue) that determines the mean phenotype and the environmental component *E* determining the phenotypic expression that maximize fitness (Θ) have a bivariate distribution with correlation *ρ* (central 95% of a bi-normal distribution with *ρ* = 0.5 is indicated by the ellipses in the lower right panel). This leads to a bivariate distribution of *U* and Θ with means *μ_u_*, variances *μ_Θ_* 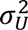 and 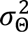, and a correlation *ρ* = *σ_uΘ_*/(*σ_u_σ_Θ_*) (top right panel). The shaded areas show the conditional probability distributions of *E* and Θ given a cue value *u* (with *ρ* = 0.5). If fitness, *W*, is a Gaussian function of the plastic phenotype value *y(u)* the optimal reaction norm as a function of cue values *u* is the same as the least squares prediction of Θ given *u, y*_opt_(*u*) = *μ*_Θ_ + 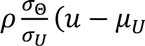 Appendix S1. Some authors refer to *U* in this context as a “proxy cue” of environmental component *Y.* However, it is sufficient to only consider *U* and Θ as two correlated components of a temporally varying environment. Blue line represents the optimal reaction norm under perfect information (*ρ* = 1) (when the ellipses collapse to a line), and green line represents the optimal reaction norm when *U* and Θ are uncorrelated (*ρ* = 0). Solid red line represents the optimal reaction norm when *ρ*=0.5 (corresponding to the drawn ellipses). Thick stippled red line is referred to in the Analysis section. Note that in Lande’s (2009) notation, *ɛ_t_* corresponds to a random value of *E* in generation *t*, and *ɛ_t_* – τ corresponds to a random *U* in the same generation.

**Figure 2.**
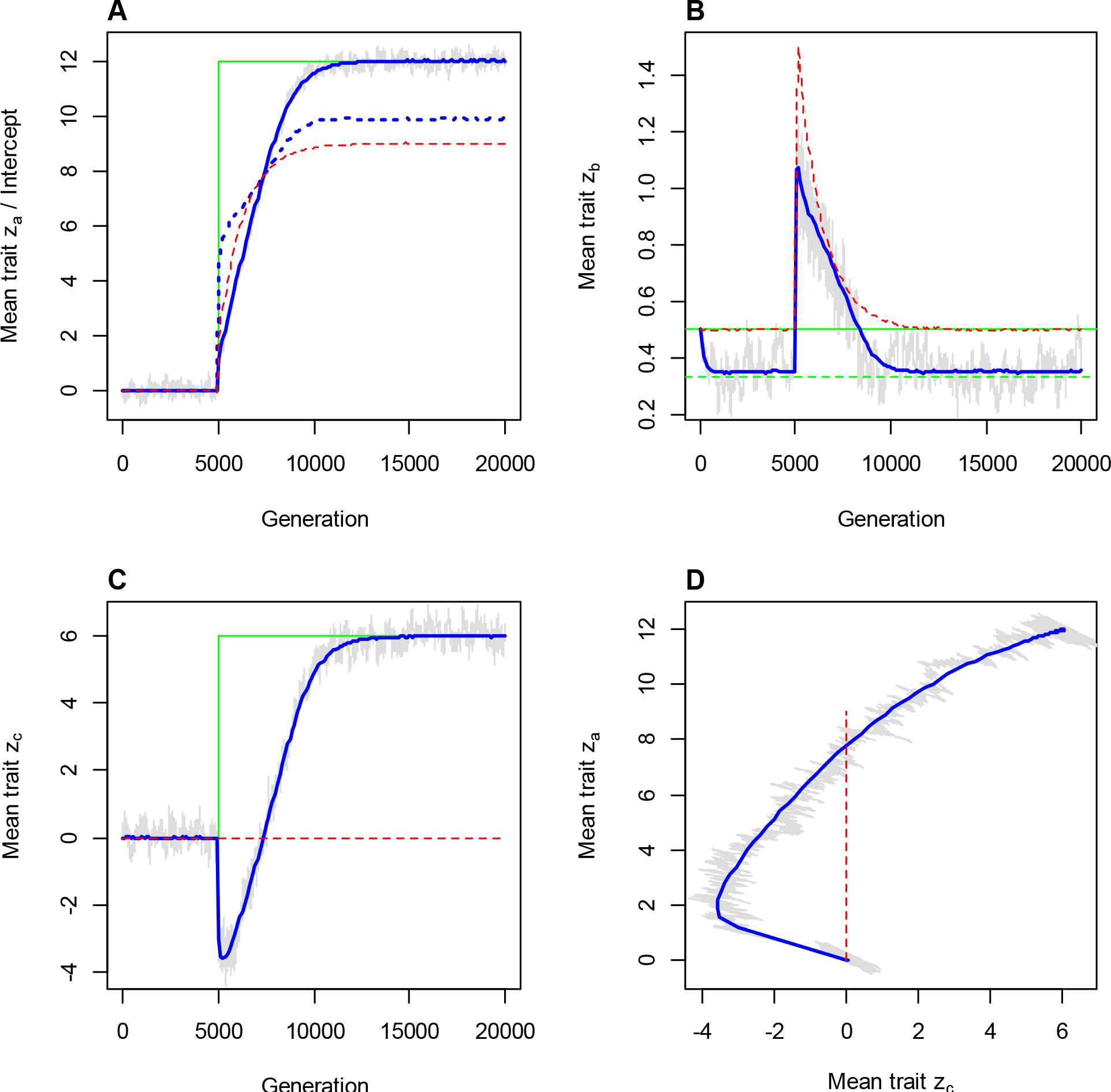
Evolution of linear reaction norms after a sudden environmental change. Panels A-C: Trajectories of the population mean trait values with a sudden environmental change at generation 5000 (see text). Panel D: Phase plane diagram showing 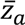 plotted against 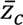 through all generations (this is the point in the cue-phenotype plane where reaction norms “tend to cross” (see Fig. 3), since the phenotypic variance-covariance matrix here is diagonal (see eqn (5)). Solid blue lines represent the three-trait model (3) and the stippled red lines represent the two-trait model (2). The trajectories were calculated as the mean of 1000 independent simulations. Grey lines show the realization of a single simulation. Solid green lines show *μ*_Θ_ (panel A), the optimal slope when reaction norm slope and intercept can be tuned independently, 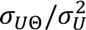 (eqn (1)) (panel B), and *μ_u_* (panel C). In panel A, the dotted blue line is the mean intercept 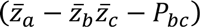 in the three-trait model for comparison with the intercept trait in the two-trait model (stippled red line). In panel B, stippled green line shows the mean slope that gives maximum expected logarithm of fitness of a random individual (Appendix S3). Parameter values in the initial environment were *μ^u^* = 0, *μ_Θ_* = 0, *σ_U_* = 2, *σ_Θ_* = 4, and 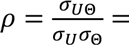 0.25. At generation 5000, *μ_u_* jumps to 6 and *μ_Θ_* jumps to 12 while the other parameters remain unchanged. Diagonal ***G*** and ***P*** matrices were used with *G_aa_ =* 0.5, *P_aa_* = *G_aa_* + 0.5, *P_bb_* = *G_bb_* = 0.045, and *ρ_cc_ = G_cc_* = 2 (three-trait model) or *P_cc_* = *G_cc_* = 0 (two-trait model). Initial mean trait values were 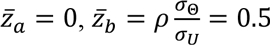 and 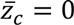

The final stage of the genetic assimilation process where the zone of canalization moves to the new mean environment is perhaps the least understood; it has been suggested that genetic drift or fitness costs of maintaining plasticity plays a part (West-Eberhard, 2003; Pigliucci *et al.*, 2006; Lande, 2009; Bateson & Gluckman, 2011), and changes in the genetic variances, covariances and genetic architecture of reaction norm components may be involved (Wagner *et al.*, 1997; Steppan *et al.*, 2002; Le Rouzic *et al.*, 2013).

One approach to quantitative genetics analysis of phenotypic plasticity (Via *et al.*, 1995; Rice, 2004) is to consider the intercept and slope of linear reaction norms as two quantitative traits in their own right (de Jong, 1990; Gavrilets & Scheiner 1993a; de Jong & Gavrilets, 2000; Tufto, 2000; Lande, 2009). More generally, reaction norms have been modeled by considering polynomial coefficients as traits (Gavrilets & Scheiner, 1993b; Scheiner, 1993). In these models, the intercept trait is defined as the value of the plastic phenotype at a reference cue designated as zero by the researcher. Lande (2009) analyzed the evolution of such a linear reaction norm, assuming a stochastic environment undergoing a sudden change (relative to the background fluctuations) in both the mean environmental cue and the phenotypic value where fitness is maximum. In his model the population responded by a rapid increase in mean reaction norm slope (plasticity), followed by a slow increase in reaction norm intercept with a concomitant decrease in plasticity. However, the genetic assimilation was not completed, as the zone of canalization could never move away from the reference cue because the covariance between reaction norm slope and intercept was assumed to remain constant. Lande (2009) argued that further reduction in phenotypic variance would take place (e.g., due to fitness costs of maintaining plasticity), but did not include any such mechanisms in his modeling.

In this paper, we argue that the two-trait model is an insufficient representation of genetic effects on linear reaction norms, and hence fails to predict critical aspects of the evolution of phenotypic plasticity and genetic assimilation. Instead we suggest modeling linear reaction norms as being composed of three traits based on the most fundamental ways that gene products may alter linear reaction norms in such a way that they remain linear. Reanalyzing the scenarios for extreme environmental change considered by Lande (2009), we show that, under the three-trait reaction norm model, genetic assimilation in the new stochastic environment becomes complete (as defined above) without changes in genetic correlations among the defined traits, genetic drift or imposing any fitness costs on maintaining plasticity. Further, we show that the evolutionary equilibrium of this three-trait linear reaction norm under random mating entails (with certain exceptions) a shallower mean reaction norm slope than the slope of the optimal individual reaction norm and the equilibrium slope of the two-trait model. Hence, maximum individual fitness does not occur at the mean trait values in the population.

We start by deriving an expression for optimal linear reaction norms as a function of environmental cues in stationary stochastic environments. We then derive our three-trait linear reaction norm model, and finally we analyze the evolutionary dynamics of this model in a quantitative genetics framework, and compare it to the dynamics of the two-trait reaction norm model analyzed by Lande (2009).

## Models

### Optimal linear reaction norms in temporally variable environments

Models for optimal adaptations in variable environments have traditionally assumed either that individuals have no information about the relevant environmental variables, or that individuals have exact information about the state of the environment (Yoshimura & Clark, 1991; Roff, 2002). Whenever the phenotype yielding highest fitness is not known exactly (i.e., the individuals do not have full information about the present and future environment), the long term success of a genotype depends not only on the expectation of fitness, but it is also adaptive to reduce the variance in mean fitness across generations (Yoshimura & Clark, 1991; Starrfelt & Kokko, 2012). Models that assume that individuals have no information about the environment have been used to explain risk-avoidance and bet-hedging strategies (den Boer, 1968; Hopper *et al.*, 2003; Starrfelt & Kokko, 2012). On the other side of the spectrum, models that predict optimal trait values as a function of environmental variables, often assume that these variables are known to the individuals without error (e.g. Stearns, 1992; Roff, 2002).

The concept that phenotypic expressions are functions of more or less informative environmental cues is well established in evolutionary ecology (Tollrian & Harvell, 1999; DeWitt & Scheiner, 2004; Stephens *et al.*, 2007; McNamara *et al.*, 2011; Gienapp *et al.*, 2014). For example, seasonal reproduction in many organisms must take place within a rather narrow time-window which often varies largely between years (Durant *et al.*, 2007; Gienapp *et al.*, 2014). Since such phenological events must often be prepared a long time in advance (due to acquiring resources, physiological developments and migration), seasonal reproduction may be influenced by rather information-poor cues such as temperature and food constituents weeks before reproductive success is determined (Berger *et al.*, 1981; Korn & Taitt, 1987; Lindstrom, 1988; Negus & Berger, 1998; Nussey *et al.*, 2005). Examples of such obviously adaptive phenotypic plasticity to more or less informative environmental cues are ubiquitous in nature (Pigliucci, 2005; Sultan, 2010; Landry & Aubin-Horth, 2014).

To derive an optimal norm of reaction to an imperfect cue, we may view the cue, *U*, and the phenotypic expression that maximize fitness, Θ, as having a joint distribution with given means, *μ_u_* and *μ_Θ_* variances, 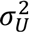 and 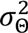 and a correlation, 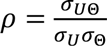 (Fig. 1). Note that we here define the cue *(U)* in a general sense as the *environmental component that* affects the phenotype, *not how* this component is perceived by the individuals (as in e.g. Tufto (2000)). Also note that *U* must not necessarily be interpreted as a proxy for another environmental component that affects fitness (e.g. Miehls *et al,* 2013), although this may be the case (see caption of Fig. 1). Hence, following McNamara *et al.* (2011) we focus on the information content in the cue *(U)* about the optimal phenotypic expression (Θ) in the given environment.

Under the assumption of no density or frequency dependence, the optimal phenotypic trait values are those that maximize the geometric mean of fitness across generations (Dempster, 1955; Caswell, 2001). This is equivalent to maximizing the expected logarithm of fitness. Hence, if fitness, *W*, is a Gaussian function (with constant width and peak value) of the phenotype value, *y*, such that (*W*(*y*)) is a quadratic function, the optimal *linear* reaction norm as a function of cue values *u* is

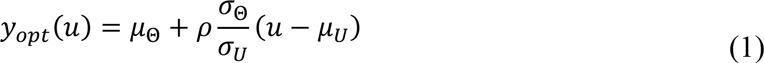

(Appendix S1). Note that, due to the quadratic fitness function In (*W*(*y*)), this is the same as the least squares prediction line of Θ as a function of cue values *u* (Battacharyya & Johnson, 1977).

This optimal individual reaction norm under imperfect information (eqn (1)) may be seen as a weighted average of the optimal phenotype under no information (*μ*_Θ_) and the optimal phenotype under perfect information (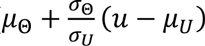), with the weight being |*ρ*| (Fig. 1). Given that *W* is a Gaussian function of *y*, this linear reaction norm is the optimal reaction norm (i.e., a non-linear reaction norm would not perform better) as long as E[Θ|*U* = *u*] is a linear function of *u*, which is the case when *U* and Θ are bi-normally distributed (chap. 7.8 Johnson and Wichern, 2007).

Optimality models of this kind have been central in the development of evolutionary ecology (Parker & Maynard Smith, 1990; Sutherland, 2005; Roff, 2010). McNamara *et al.* (2011) analyzed the general optimal linear reaction norm given by eqn (1) in terms of optimal phenology under environmental change. Ergon (2007) used a similar approach to analyze optimal trade-offs between pre-breeding survival, onset of seasonal reproduction and reproductive success in fluctuating multivoltine species.

### Quantitative genetics models for linear reaction norms – two vs. three traits

The optimal linear reaction norm given by eqn (1) says nothing about the selection process and does not consider genetic constraints. In the following we will consider a quantitative genetics model for linear reaction norms, assuming phenotypic responses to an interval-scaled cue with an arbitrary zero point (Houle *et al*, 2011).

In quantitative genetics models for the evolution of phenotypic plasticity, it is common to consider the intercept (*α*) and slope (*β*) of the reaction norm as two traits (e.g. de Jong, 1990; Gavrilets & Scheiner, 1993a; de Jong & Gavrilets, 2000; Tufto, 2000; Lande, 2009; Scheiner, 2013). I.e., the plastic phenotype is modeled as a function of an environmental cue *u* on the form

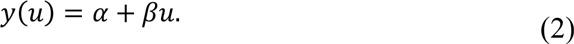

In this two-trait model, the intercept trait *α* is the phenotypic expression for the cue-value designated as zero. Lande (2009) assumed that minimum phenotypic variation occurred in the mean environment that the population had been adapted to, and hence defined the cue to have its zero point in this reference environment. He then used this reaction norm model (eqn (2)) in a quantitative genetics analysis of adaptations to a sudden extreme change in the mean environment when the reference environment remained unchanged.

We will here analyze a more general linear reaction norm model based on the three most fundamental ways that genetic effects can alter a linear reaction norm in such a way that it remains linear; (i) a change along the plastic phenotype axis, (ii) a change in slope (cue sensitivity), and (iii) a change in the reaction norm along the cue axis. This leads us to consider a linear reaction model on the form

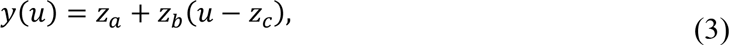

where *z_a_*, *z_b_*, and *z_c_* are considered as (latent) traits. A particular genetic effect may of course affect more than one of these traits, but any genetic effect on a linear reaction norm can be decomposed into these three components. Obviously, shifting a linear reaction norm along the cue-axis (a change in *z_c_*) may have exactly the same effect on the reaction norm as shifting it along the y-axis (a change in *z_a_*). By rearranging the reaction norm model (3) as *y*(*u*) = *α z_b_u* where *α* = *z_a_ – z_b_z_c_*, we see that increasing *z_a_* by one unit has the same effect on *y*(*u*) as decreasing *z_c_* by 1/*z_b_* units. However, traits *z_a_* and *z_c_* still represent very different genetic effects within the organisms. Trait *z_c_* may be thought of as representing genetic effects on “perception” of the environmental cue in a general sense. For example, variation in *z_c_* may represent genetic effects affecting the sensory apparatus in such a way that different genotypes perceive the same environmental cue as different, but cue perception may not necessarily involve a sensory apparatus (see Discussion). Note that the intercept (*z_a_ – z_b_z_c_*) depends on the chosen zero-point of the interval scaled cue, while trait *z_a_* represents genetic effects that are invariant to which environment that has been designated (by the researcher) to have cue value zero. Variation in trait *z_a_* may thus represent variation in gene products for which both the production of these gene products and their effect on *y(u)* are independent of the cue. Finally, trait *z_b_* (reaction norm slope) represents variation in gene products that affect the sensitivity of the plastic phenotype *y(u)* to the cue. With this parameterization of the reaction norm (eqn (3)), *z_c_* may be referred to as a “cue reference trait” although we do not suggest that there is necessarily a “template” of a specific environment that is stored genetically in the organisms; what is essential is the types of genetic variation that is represented by the three traits in the model. Note that it is only when assuming a linear reaction norm that genetic effects on cue “perception” can lead to the same change in the reaction norm as genetic effects on the environment independent component of the plastic phenotype (*z_a_*); this will not be the case in a non-linear reaction norm model.

The two-trait model (eqn (2)) is a special case of the more general three-trait model (eqn (3)) where *z_c_* is fixed to zero. Reaction norm slope is considered as a trait in both models (i.e., *β* = *z_b_)*, but for clarity we have used separate notations in the two models.

## Analysis

### Basic properties of the reaction norm models

As already noted, an obvious difference between the two-trait (eqn (2)) and the three-trait (eqn (3)) reaction norm models is that the two-trait model implies a one-to-one correspondence between genotypes and reaction norms, whereas the three-trait model implies that one reaction norm can represent many genotypes. As we will see below, linear reaction norms in a population will evolve very differently and reach different equilibria when we consider the reaction norm to result from three traits rather than two traits.

An essential difference between the two-trait and the three-trait reaction norm models relates to constraints in the evolution of the covariance between reaction norm intercept and slope in the population. To see this, it is elucidating to consider a particular representation of this covariance, *u*_0_, defined as the cue value for which phenotypic variance is at a minimum and where the covariance between the plastic phenotypic value *y*(*u*) and reaction norm slope is zero (the “zone of canalization” at the population level is centered around *u*_0_). Given a phenotypic covariance between intercept and slope (*P_αβ_*) and a variance in reaction norm slope (*P_ββ_*), this cue value is

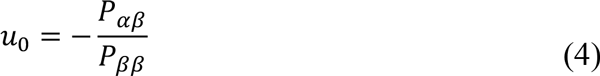

(Appendix S2).

From eqn (4) we see that, in the two-trait model (2), where reaction norm intercept (*α*) and slope (*β*) are considered as traits, *u*_0_ is independent of the trait means, and directional selection on any of the traits will not affect *u*_0_ unless the selection also changes the variance of the slope or covariance of the traits.

On the other hand, in the three-trait model (3), the covariance between intercept and slope depends on the mean traits 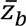 and 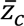 Under the assumption of normal traits, *u*_0_ then becomes

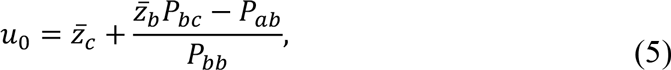

where *P_bc_,P_ab_ and P_bb_* are the elements of the phenotypic variance-covariance matrix indicated by the subscripts (Appendix S2). Thus, under the three-trait model (3), *u*_0_ may respond directly to directional selection on both trait *z_b_* (if *P_bc_* ≠ 0) and trait *z_c_.* If trait *z_b_* is independent of trait *z_a_* and *z_c_* (i.e., *P_bc_* = *P_ab_* = 0), *u_0_* becomes 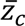. Note also that *u*_0_ is independent of *P_ac_.*

Lande (2009) defined the cue *u*(*ɛ_t-τ_* in his model) to have its zero-point at *u*_0_ as a “reference environment”. Hence, one could define the two-trait model analyzed by Lande (2009) for any arbitrary interval scaled cue variable as *y*(*u*) = *α^′^* + *β* (*u –u*_0_) where the genetic correlation between the traits *α^′^* and *β* is by necessity zero since *u*_0_ is defined by *cov* (*y*(*u*_0_),*β*) = *cov*(*α^′^,β*) = 0 (Appendix S2; see also last paragraph on page 1438 in Lande (2009)). This model is structurally similar to our three-trait model except that the “reference environment” in our model is considered as an individual trait, *z_c_* (reflecting individual variation in cue “perception”), which is exposed to selection. Unlike in Lande’s (2009) model, where the definition of trait *α^′^* depends on *u*_0_, there are no constraints on the phenotypic or genotypic covariances in our three-trait model (other than that the covariance matrix must be positive-definite). The two-trait model of Lande (2009) can only evolve in the same way as the three-trait model if *u*_0_ is treated as the mean of an individual trait with variance different from zero. Hence, the three-trait quantitative genetics model and Lande’s (2009) two-trait model are not alternative parameterizations of the same model. Lande’s (2009) two-trait model is a constrained version of our more general three-trait model with the trait *z_c_* fixed to *u*_0_, which requires that *P_cc_* = *P_ac_* = *P_bc_* = 0 as well as *P_ab_* = 0 (*P_ab_* = 0 is only required to maintain the same definition of *z_a_* and *α^′^* and to give 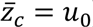). We will later show that expected *u*_0_ at equilibrium in the three-trait model always becomes *μ_U_*.

### Evolution of linear reaction norms

Environmental change may lead to changes in any of the parameters of the joint distribution of cue (U) and the best possible phenotype (Θ) (cf., eqn (1) and Fig. 1). Any such change will impose directional selection on the individual traits defining the reaction norm, and the evolutionary response to this selection will depend on the additive genetic variances and covariances of these traits. We will here compare the evolution of linear reaction norms based on the three-trait model (eqn (3)) and the more constrained two-trait model (eqn (2)) analyzed in detail by Lande (2009). Specifically, we will analyze the transient and asymptotic evolution of the reaction norm distribution after a sudden and extreme concomitant change in both *μ_U_* and *μ_Θ_*, while 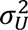, 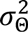 and *σ*_UΘ_ remain unchanged. We assume that all individuals in each generation experience the same environment, and that the environments in subsequent generations are independent (as also in Lande’s (2009) analysis). Following Lande (2009) we also assume that trait variances and covariances remain constant under selection. Although this may be a particularly unrealistic assumption (Steppan *et al*, 2002), it serves the purpose of examining how reaction norms can evolve through changes in trait means only.

### Quantitative genetics – modeling

Assuming that the individual traits of the reaction norm (3) have a multi-normal distribution with a constant variance-covariance matrix in a population with discrete generations, the fundamental equation describing the change in the population mean of the traits from a generation *t* to the next,

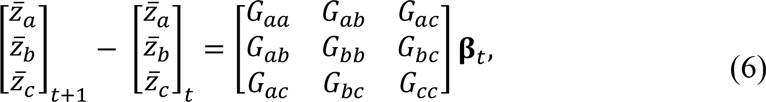

is the product of the additive genetic variance-covariance matrix for the traits, ***G***, and the selection gradient **β***_t_*. Here, **β***_t_* is the sensitivity of the logarithm of population mean fitness to changes in each of the mean trait values (Lande, 1979, Lande & Arnold, 1983),

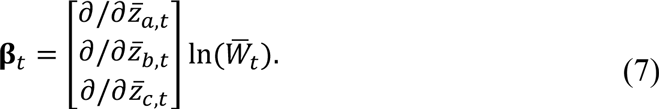

We will assume a Gaussian fitness function with width *ω* and peak value *W_max_*, and that all individuals experience the same environment in any generation.

A random individual in generation *t* has phenotype *y_t_*(*u_t_*) = *Z_a,t_* + *Z_b,t_*(*u_t_* – *Z_c,t_*), where the traits [*Z_a,t_,Z_b,t_,Z_c,t_*] are drawn from a multi-normal distribution with mean 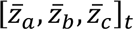 and phenotypic covariance matrix ***P.*** When the phenotypic expression that maximizes fitness in that generation is *θ_t_*, this individual will have fitness

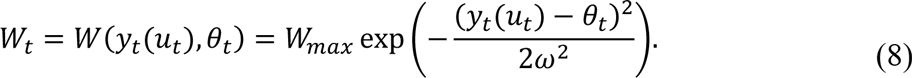

To find an analytical expression of the selection gradient (7), a common approach (Lande and Arnold 1983, Lande 2009) would be to first find the population mean fitness by integrating over the phenotype distribution, *p*(*y_t_*(*u_t_*)),

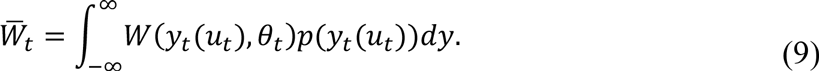

However, because *p*(*y_t_*(*u_t_*)) is not normal as it involves the product of the two normally distributed traits *z_b,t_* and *z_b,t_*, it is not straightforward to solve this integral analytically. Indeed, it seems that an exact analytical expression for the selection gradient (7) does not exist. We therefore initially based our analysis on simulations of the evolutionary process (6), where the selection gradient (7) is computed numerically by simulating a population of 10,000 individuals at each generation (see Appendix S5 for R code). These simulations are accompanied by (and compared to) mathematical analyses presented in Appendix S3 and Appendix S4.

In the simulation results presented in Fig. 2, we used the same parameter values as in Lande’s (2009) analysis of the two-trait model except that we, for convenience, used a somewhat less extreme sudden change in the environment, with a change in *μ_U_* and *μ_Θ_* of 3 (instead of 5) standard deviations of the background fluctuations (*σ_U_* and *σ_Θ_* of eqn (1)). As Lande (2009) we used a diagonal **G**-matrix and sat *G_cc_* to half the cue variance (three-trait model) or zero (two-trait model). For simplicity, in the simulations we also assumed that only trait *z_a_* had a non-additive residual component with variance 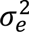 such that 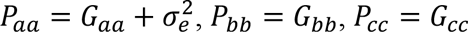, and *P_ab_* = *P_ac_ P_bc_* = 0. The two-trait model is obtain simply by setting also *P_cc_* = 0 and 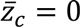.

### Quantitative genetics – results

The simulations show that immediately after the sudden environmental change, there is a rapid increase in reaction norm slope (Fig. 2B), while 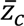 (Fig. 2C) swings back in the opposite direction of the change in mean cue *μ_U_* (i.e., away from the new optimum). This phase of the adaptation may be characterized as a “ state of alarm”, where it becomes adaptive to exaggerate the perception of the environmental change. As 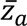 moves towards the new optimum (Fig. 2A), the reaction norm slope 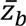 is reduced and 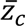 turns towards the new optimum. Eventually, 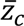 stabilizes around *μ_U_* and 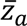 stabilizes around *μ*_Θ_ (Fig. 2D), in accordance with the theoretical results in Appendix S3 (see Appendix S4 for detailed numerical results). Note that with *P_ab_* = *P_bc_* = 0 (as in the simulations), the theoretical equilibrium mean values 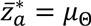 and 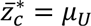, are independent of the variances and covariance of *U* and Θ. In Appendix S4 we conjecture that the equilibrium mean traits 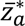 and 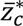 in general (for *P_ab_* ≠ 0 and *P_bc_* ≠ 0) are affected by 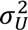, 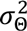 and *σ_UΘ_*, but then only indirectly through 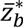.

Since we used a diagonal phenotypic variance-covariance matrix (**P**) in the simulations, the cue value *u*_0_ that yields minimum phenotypic variance (eqn (5)) equals 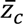, which stabilizes around the theoretical equilibrium *μ_U_* (Fig. 2C; Appendix S3). Hence, in this case, equilibrium *u*_0_ becomes 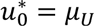. As shown both by simulations (Supporting Figs S1-S3) and theoretical considerations (Appendix S4), this property (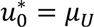) holds also when ***P*** is not diagonal – i.e., at equilibrium, phenotypic variance is *always* minimum in the mean environment. As a result, the three-trait model leads to complete genetic assimilation in the sense that the population level zone of canalization (represented by *u_0_)* evolves to the mean environment regardless of what this mean is. In contrast, in the two-trait model, *u*_0_ does not evolve in response to changes in the trait means and the phenotypic variance can only be minimized when the mean environment equals –*P_αβ_* / *P_ββ_* (see eqn (4)). This contrast in the asymptotic state of the systems obtained from the two alternative reaction norm models is illustrated in Fig. 3, and Fig. 4 shows the trajectories of phenotypic variation and difference between *u*_0_ and *μ_u_* in the simulated scenario presented in Fig. 2. Supporting Figures S4 and S5 show simulation results for a scenario where there is no environmental variation before and after the sudden environmental change (more similar to classic examples of genetic assimilation).

**Figure 3.**
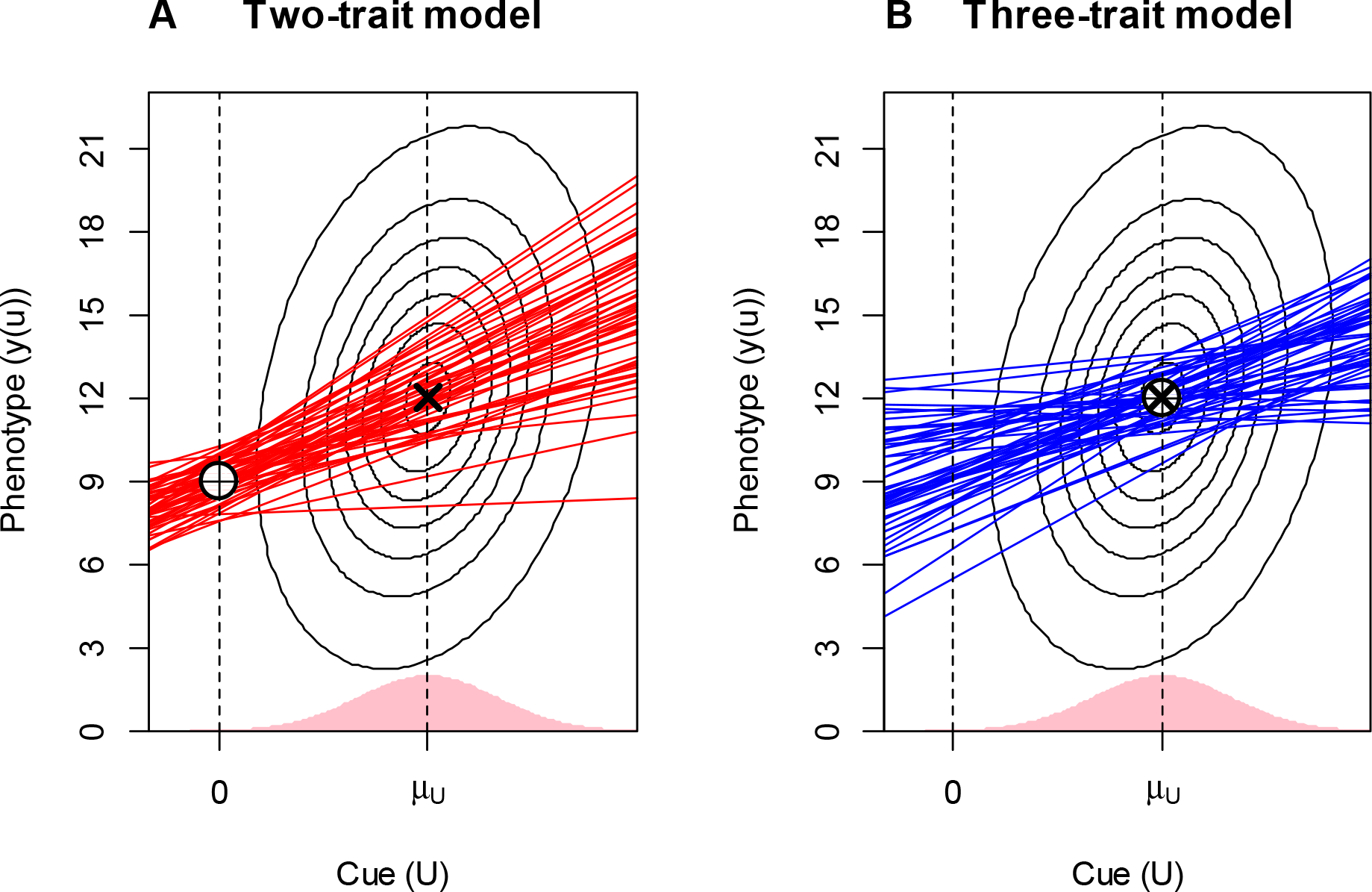
Reaction norm distribution when the populations have reached a stationary dynamics in the two-trait model (A) and the three-trait model (B) under the scenario presented in Fig. 2. The distribution of the environmental cue *(U)* in the new environment is indicated by the shaded areas on the x-axes, and the central 95% of the joint distribution of *U* and Θ is shown with the ellipses with an ‘ **×**’ at the mean. For each model, 50 random reaction norms (genotypes) are plotted. In the two-trait model, the cue value *u*_0_ where phenotypic variation is minimal will always be at zero when reaction norm slope and intercept are independent (indicated with a white, crossed, symbol plotted at the mean plastic phenotype for this cue value). In contrast, in the three-trait model genetic assimilation becomes complete and *u*_0_ moves to with a mean plastic phenotype at *u*_0_ moves to *μ_U_* with a mean plastic phenotype at *μ_Θ_*

Interestingly, as seen in Fig. 2B, the mean reaction norm slope 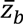 in the three-trait model stabilizes at a lower level than the optimal slope yielding the highest expected fitness of an individual, 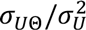 (see eqn (1)), which is also the equilibrium mean slope in the two-trait model (Gavrilets & Scheiner, 1993a; Lande, 2009). Intuitively, this is because the optimal value of trait *z_b_* of an individual depends on the value of trait *z_c_* that this individual possesses, which is stochastic. Under the assumption that *P*_ab_ = 0, an approximate mean slope value is found as 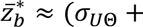 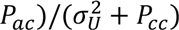 (Appendix S4 and eqn (10) below),which is close to the stationary mean in the simulations (Fig. 2). For comparison, the equilibrium mean traits in the two-trait model become 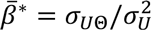 and 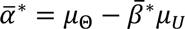 (Gavrilets & Scheiner, 1993a; Lande, 2009). Note that the denominator in the approximate expression for 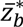 is the variance (*U-Z_c_*), and not the variance of the cue *U* alone as in the expression for 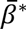 in the two-trait model; i.e., genetic variance in the perception trait *z_c_* inflates the variance of the perceived cue *U – Z_c_*). Hence, if 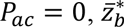 is always lower than the optimal slope in eqn (1) unless *P*_cc_ = 0 (which gives the two-trait reaction norm model). This is indicated by a stippled reaction norm in Fig. 1.

As seen in Fig. 2B the asymptotic mean 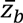 in the simulations (where is *P_ac_* = 0) is close to but somewhat larger than the approximation 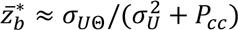. This discrepancy is further analyzed in Appendix S4. As shown there, the equilibrium mean reaction norm slope 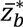 can be approximated analytically if we assume that the plastic phenotype *y(u)* has a normal distribution, which is very nearly the case with the parameter values in our simulations in Fig. 2. The integral (9) then has an analytical solution, and as a result an approximate equilibrium slope 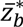 can be found numerically from the equation (assuming *P_ab_* = *P_bc_* = 0)

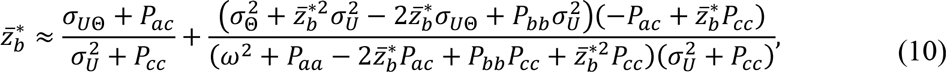

where the large values of 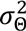 and especially *ω*^2^ used in the simulations make the second term positive but small compared to the first term (see Appendix S4 for detailed numerical results).

Another reason for the discrepancy between the asymptotic mean 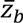 in the simulations and the approximation 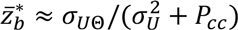 is that when the population under directional selection based on eqn (6) evolves towards a stationary state, the mean traits will fluctuate around the equilibrium because of the influence from the random inputs *u_t_* and *Θ_t_* (as seen in Fig. 2). In stationarity this leads to 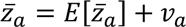 etc. (where 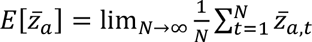 and *E*[*ν_a_*] = 0 etc.), and, as shown in Appendix S4, the variances and covariances of *v_a_, v_b_* and *v_c_* then enter into eqn (10). Note that we assume that *u_t_* and *θ_t_* have zero autocorrelation, such that the covariances between the mean reaction norm parameters and the environment caused by adaptive tracking (Tufto, 2015) are zero.

Because the reaction norm slope 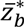 is influenced by the phenotypic variance of the cue reference trait *z_c_* (and its covariance with the other traits; eqn (10)), and hence deviates from the slope that maximizes fitness (eqn (1)), the expected fitness at equilibrium will be lower than the expected fitness of the optimal individual reaction norm in eqn (1) (Fig. 5, lower right panel). As a consequence a proportion of the population will have a higher expected fitness than an individual with mean trait values. Nevertheless, mean fitness in the population after the environmental change stabilizes around a higher level in the three-trait model than in the two-trait model (Fig. 5, left panels), despite a lower expected fitness at mean trait values (right panels). The reason for this is that the three-trait model gives a lower phenotypic variance in the new environment (Fig. 4A). Mean fitness in the two-trait model thus stabilizes around the optimum *only when* the mean cue is zero because phenotypic variance will not be minimized in other environments (Fig. 5, left panels).

**Figure 4.**
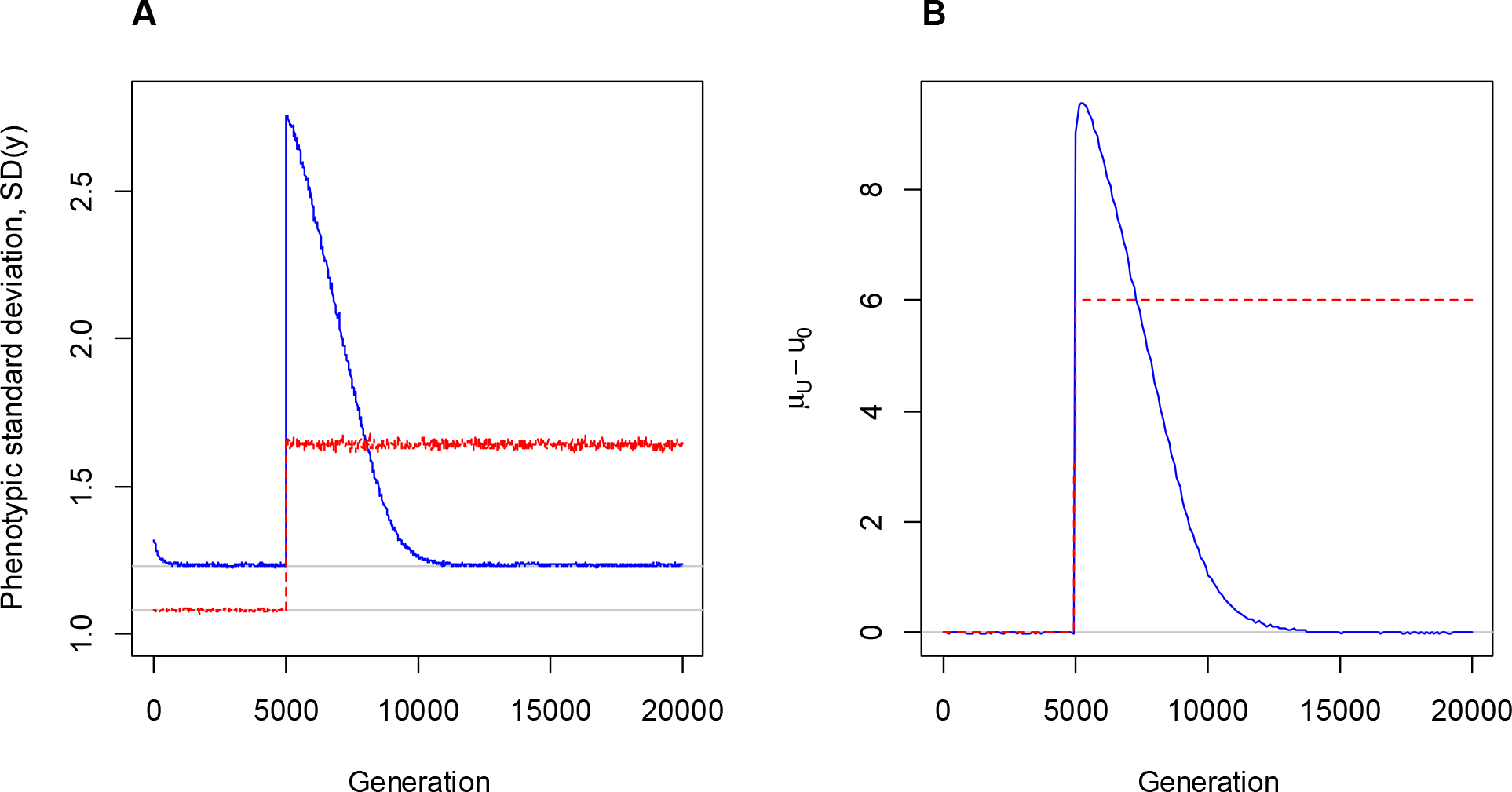
Phenotypic standard deviation, (**A**), and the distance between the mean environment and the zone of canalization, *μ*_u_ – *u*_0_ (**B**), in the simulations presented in Fig. 2. Blue solid lines represent the three-trait model, while the red stippled lines represent the two-trait model. Horizontal grey lines are drawn at the mean values of the last 3000 generations prior to the sudden environmental change at generation 5000. Lines show the mean of 1000 independent simulations plotted at every 100^th^ generation.

**Figure 5.**
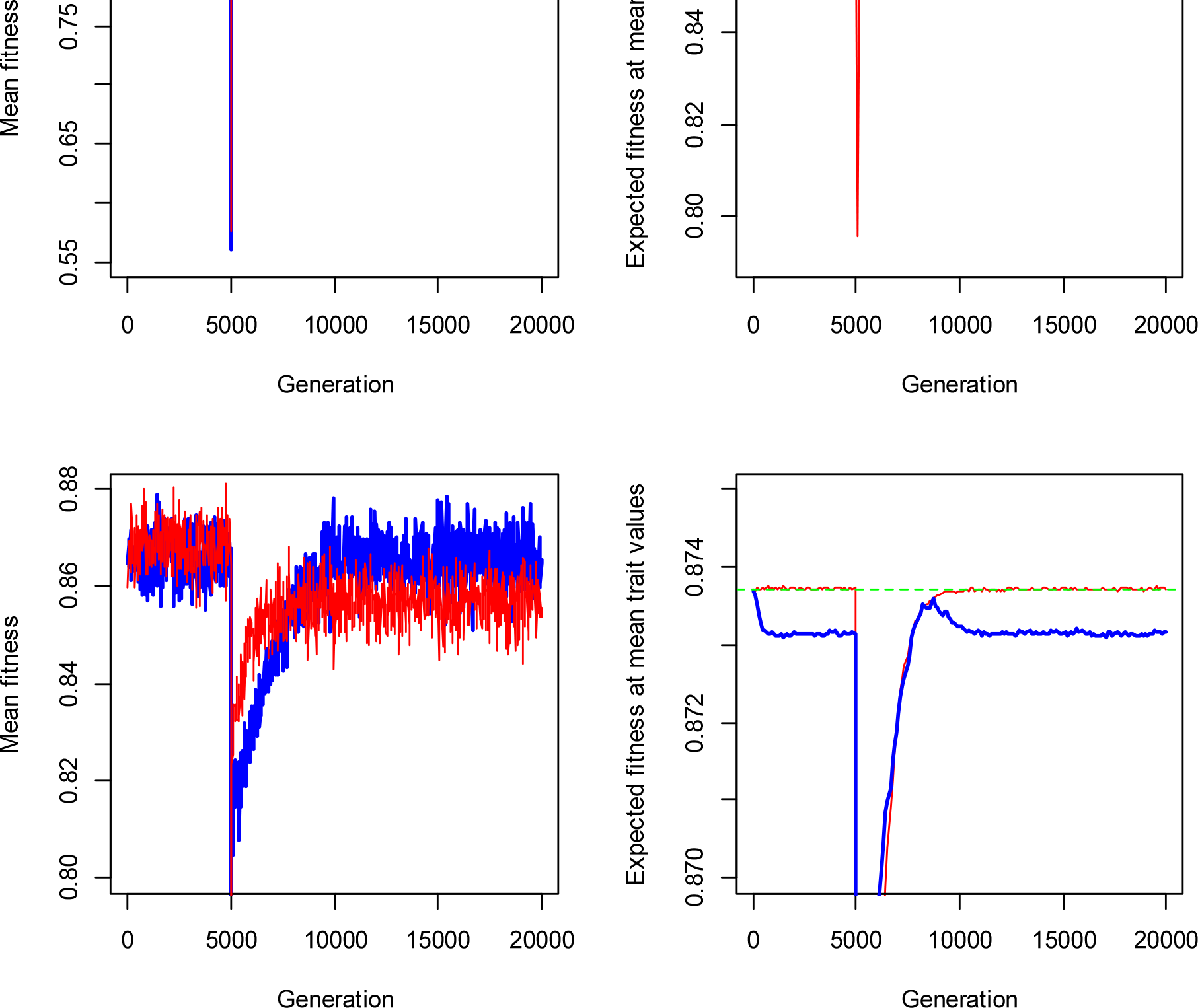
Fitness trajectories in the simulation example (Fig. 2). Left panels: Population mean fitness relative to maximum fitness (*W_max_*). Right panels: Expected fitness at the mean trait values. The lower panels show the same values plotted with a narrower range on the y-axis. Thick blue line represents the three-trait model and the thin red line represents the two-trait model. Horizontal stippled green line shows the fitness of the optimal reaction norm (1). Only every 20^th^ generation is plotted in the left panels and every 100^th^ generation is plotted in the right panels. Plotted values are the mean of the same 1000 independent simulations used for Fig. 2.

## Discussion

Quantitative genetics models are theoretical models for the joint evolution of population means of quantitative individual phenotypic traits, where the researchers define traits that they find most meaningful in the context they are studied. In quantitative genetics models of reaction norms where a plastic phenotype is modeled as a linear function of an interval scaled environmental cue, the reaction norm intercept and slope are often considered as individual traits subjected to selection (Gavrilets & Scheiner, 1993b; Scheiner, 1993; de Jong & Gavrilets, 2000; Tufto, 2000; Lande, 2009; Scheiner, 2013; Tufto, 2015). The intercept of such a reaction norm (i.e., the reaction norm value at cue value zero) is often not very biologically meaningful since this trait, as well as its variance and covariance with other traits, depend on the defined zero-point, or “reference cue”, of the (arbitrary) interval scaled cue variable. One may, however, as in Lande (2009), define the zero-point of the cue to be the mean cue value which the population is adapted to. This ensures that the variance of the plastic phenotype is minimized in the mean environment, which is theoretically plausible (Bürger, 2000; Lande, 2009; Le Rouzic *et al*, 2013), but it is not clear how this “reference cue” may evolve (in Lande’s (2009) analysis it is assumed to remain constant; see however de Jong & Gavrilets 2000).

We have here suggested that the “reference cue” can be considered as an individual trait that reflects genetic variation in cue “perception” in a general sense, and hence considered a linear reaction norm on the form *y(u) = z_a_ + z_b_*(*u – z_c_*). In this model, the biological meaning of all the traits, and their variances and covariances, is not modified when redefining the zero-point of the cue variable *u* (which is not the case for the intercept *α = z_a_ + z_b_z_c_, var(α)* and *cov (α, z_b_*)). The three traits in this model reflect three fundamentally different genetic effects on linear reaction norms. While *z_b_* represents genetic effects on cue sensitivity, *z_c_* reflects genetic effects on cue “perception” (in the general sense discussed below) and has the same scale as the environmental cue, and *z_a_* represents genetic effects that are both independent of the cue value and invariant to its defined zero-point (the latter is not the case for the intercept). These structural differences in the reaction norm models matter for the equilibrium mean reaction norms (and distributions), because the traits do not have independent effects on the plastic phenotype (*y (u)*) (note the product *z_b_z_c_* in the three-trait model).

In our analysis of the three-trait model, we have shown that the cue value where variance of the plastic phenotype is minimized (where reaction norms “tend to cross”; *u*_0_) always evolves to equal the mean environment at equilibrium. This occurs without assuming any cost of maintaining plasticity (DeWitt *et al*, 1998; West-Eberhard, 2003; Pigliucci *et al*, 2006; Lande 2009; Bateson & Gluckman 2011; Svennungsen *et al*, 2011), or any change in the variances or covariances of our defined traits (de Jong & Gavrilets, 2000). Even though *u*_0_ may be interpreted as ‘–(intercept, slope)/*var*(slope), *u*_0_ is biologically more meaningful than the covariance between reaction norm slope and a somewhat arbitrarily defined intercept trait. Note that *u*_0_ is a population level parameter that does not depend on any quantitative genetics model for the linear reaction norm, and which can easily be estimated (as discussed below). Further, our analysis also demonstrate that the equilibrium mean reaction norm slope in the three-trait model will deviate from the optimal slope yielding the highest expected fitness of a hypothetical individual that can tune reaction norm intercept and slope accurately and independently (eqn (1)), which is also the equilibrium mean slope of the two-trait model (Gavrilets & Scheiner 1993a; Lande, 2009). At least when there is weak correlation between *z_a_* and *z_c_* (i.e., *P_ac_* is sufficiently small), the equilibrium mean slope will be lower than the optimal individual slope. Intuitively, this is because the optimal slope is lower when the cue reference trait of a random individual, in addition to the environmental cue, is stochastic due to random mating. As a consequence, maximum expected fitness does not occur at the mean trait values in the population.

In the three-trait model, phenotypic variance in a given environment increases with both 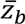 and the distance between 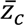 and the environmental cue *(u)*, at least when the traits are independent (see eqn S4-3 in Appendix S4), whereas in the two-trait model, phenotypic variance is independent of the trait means (Fig. 4). In our simulations, after the sudden environmental change, there is a rapid initial increase in both 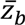 and the distance between 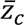 and the new mean cue value (i.e., 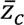 initially evolves rapidly in the *opposite* direction of the change in the environmental cue, such that the perception of the environmental change is exaggerated). Hence, due to the positively interacting (epistatic) effects of 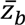 and 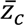 on the plastic phenotype *y*(*u*) this efficiently increases phenotypic variance in the new environment which enhances the evolvability of the plastic phenotypic character and acts to restore population mean fitness (see Figs 2 and 5). The subsequent process of assimilation where by reaction norm slope 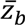 is reduced, 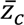 moves towards the mean cue value, and 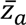 evolves towards mean Θ, is a much slower process.

### Genetic effects on linear reaction norms

Although a shift in the reaction norm along the cue-axis (through trait *z_c_*) can have exactly the same effect on the individual linear reaction norm as a shift along the phenotype-axis (through trait *z_a_*), the genetic bases for these effects are fundamentally different, and, as explained above, changes in the means of these two traits have different effects on the population. It also seems obvious that there will often be genetic variation on both these traits.

Phenotypic plasticity involves complex pathways, at both organismal and cell levels, from perception of environmental cues and physiological transduction to phenotypic expression (reviewed in Sultan & Stearns, 2005). Depending on the type of organism and the nature of the phenotypic characters and the environmental cues, these pathways may, to varying degrees, involve sensory systems, neuroendocrine and metabolic systems, cellular reception, gene regulation networks, and other developmental, physiological and behavioral processes. Environmental conditions may directly affect any of these systems and processes, not just the sensory systems (e.g., temperature may directly affect metabolism and gene regulation in ectothermic organisms (Gillooly *et al*, 2002; Ellers *et al.* 2008), and various processes may be affected by food constituents (Sanders *et al*, 1981; Meek *et al*, 1995; Krol *et al*, 2012) and nutritional state (Lõmus & Sundström, 2004; Rui, 2013; Mueller *et al*, 2015)). Genetic variation in upstream (i.e., close to the cue perception) regulatory processes, which may involve cue activation thresholds for transduction elements, may affect the way the environment is “perceived” (in a general sense) by the organism, and hence the cue reference trait (trait *z_c_*) in our model. Genetic variation in downstream processes close to the phenotypic expression of quantitative characters, on the other hand, may affect the degree of up/down regulation in response to given levels (and types) of transduction elements and hence the slope of linear reaction norms (trait *z_b_* in our model). Finally, some genetic variation may have the same additive effect on the phenotype irrespective of the environmental cue (trait *z_a_* in our model). The importance of differentiating between these three traits may be better appreciated when considering the effects of the mean traits on the population; A change in 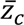 will change the cue value at which different genotypic reaction norms tend to cross *(u_0_)*, whereas a change 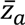 will not.

While there is ample evidence for widespread genetic variation for reaction norms in natural populations (Falconer & Mackay, 1996; Sultan & Stearns, 2005; Sengupta *et al*, 2016), there are not many examples where the full pathway of phenotypic plasticity from cue perception to phenotype expression is known in great detail (Sultan, 2010; Morris & Rogers, 2014), and even less is known about the genetic variation of the different elements of these pathways. It seems, however, obvious that there may be substantial genotypic variation in perception of environmental cues (i.e., variation in trait *z_c_* in our model). Examples indicating genetic variation in environmental perception include substantial among-population variation in the signal transduction pathway of induced plant defense in *Arabidopsis thaliana* (Kliebenstein, *et al*, 2002), and individual variation in systemic stress responses has likely components of individual variation in what is perceived as stressful (Hoffmann & Parsons, 1991; Badyaev, 2005; Dingemanse, *et al*, 2010). There is also considerable variation and “fine tuning” in light (and shading) perception systems involving phytochromes that are sensitive to different wave lengths in plants (Smith, 1990; 1995; Schlichting & Smith, 2002).

### Predictions and empirical evaluations

Parameters in a reaction norm function considered as quantitative traits are always latent in the sense that one cannot measure their phenotypic value by a single measurement of an individual (except for traits that are defined for a particular environment, such as an intercept). While one may estimate reaction norm intercept and slope from multiple measurement of the same genotype or related individuals with known genealogy (Nussey *et al*, 2007; Martin *et al*, 2011), such data alone does not provide enough information to separate the traits *z_a_* and *z_c_* (from a statistical point of view, the three-trait model fitted to such data is over-parameterized, which may be one of the reasons it has not previously been considered; note however that the three-trait model predicts a different phenotypic distribution than the two-trait model due to the product *z_b_z_c_*). Nevertheless, if one have a detailed understanding of the physiological (or developmental) mechanisms of the plastic response one may still be able to estimate meaningful reaction norm traits beyond a phenomenological ‘intercept’ and ‘slope’, including traits associated with cue perception (trait *z_c_*). Time-series data from selection experiments may also provide information about the genetic architecture of the reaction norms (Fuller *et al*, 2005).

The cue value that gives minimum phenotypic variation in the population (*u*_0_), may be estimated by fitting data on genotype specific phenotypic measurements to mixed-effects linear models with random individual slopes and intercepts (Martin *et al*, 2011; Bates *et al*, 2015), or from a random regression “animal model” building on a known relatedness among individuals (Nussey *et al*, 2007). Our three-trait quantitative genetics model gives certain predictions about the evolution of *u*_0_ under environmental change. Our analysis shows that the mean cue reference trait (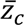), and hence *u*_0_ (eqn (5)), will respond rapidly to changes in the mean environment (provided sufficient additive genetic variation). Whenever there is selection for increased plasticity (i.e., selection for higher 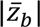), it also becomes adaptive to exaggerate the perception of the environmental change, and *u*_0_ will swing away in the *opposite* direction of the change in the mean cue during a “phase of alarm” (see Fig. 2). Later, *u*_0_ will move towards, and eventually fluctuate around, the new cue value. In contrast, under the two-trait model *u*_0_ will not change in response to changes in the mean cue values.

### Future directions

In this paper we have made a number of simplistic, but quite standard, assumptions, including interval scaled cues and phenotypes, Gaussian fitness with constant width and peak value, lack of density and frequency dependence, random mating, discrete generations where all individuals are exposed to the same environment (e.g. no spatial heterogeneity), and uncorrelated environments from one generation to the next. These assumptions may be modified or relaxed in future developments. In particular, the two-trait model has been used in theoretical studies involving within-generation heterogeneity (de Jong & Gavrilets, 2000; Tufto, 2000; Scheiner, 2013; Tufto, 2015). We suggest that these studies may be developed by including a cue reference trait in the linear reaction norms (our three-trait model). The models may also be modified by incorporating different reaction norm shapes. Notably, de Jong and Gavrilets (2000) allowed the genetic covariance between reaction norm intercept and slope, as well as their variances, to evolve through selection on allelic pleiotropy. It would be interesting to repeat their approach on our three-trait model to investigate the relative contributions (and synergies) of the evolution of trait means and trait variances and covariances.

Several authors have assumed flexible polynomial reaction norms with the polynomial coefficients considered as traits (Gavrilets & Scheiner 1993a, b; Scheiner, 1993, Via *et al*, 1995). We suggest that such rather phenomenological non-linear reaction norm models may be modified by considering the slope and perception traits of the three-trait model as themselves dependent on the environment, which may result in a polynomial of *(u —* z_c_); Considering trait *z_b_* as a linear function of *(u — z_c_*) results in a reaction norm that is a second order polynomial of *(u — z_c_*), etc. Note that in nonlinear reaction norms, unlike linear ones, a change in the perception trait(s) will never have the same effect on the genotypic reaction norm as a change in elevation trait (the component of the plastic phenotype independent of the environment).

Regardless of the reaction norm shape, we argue that it is essential to distinguish between genetic variation in how the environmental cues are perceived from other genetic variation affecting the reaction norm distribution in the population. We suggest that future developmental and behavioral studies pay more attention to genetic variation in environment perception and transduction, and that the contributions of such genetic variation to phenotypic variation in natural environments are evaluated.

## Acknowledgements

We thank Arnaud Le Rouzic, Richard Gomulkiewicz, Samuel M. Scheiner, Thomas F. Hansen and Øistein H. Holen for comments on earlier versions of the manuscript; we greatly appreciate their input and advice but may still disagree on some issues. This work made use of the Abel computing cluster, owned by the University of Oslo and the Norwegian meta-centre for High Performance Computing (NOTUR). We are thankful for technical support from the Research Computing Services group at USIT, University of Oslo.

## Supporting information

Additional Supporting Information may be found in the online version of this article:

**Supporting Figures S1 to S5.**

**Appendix S1** Optimal reaction norms.

**Appendix S2** Cue value *u*_0_ where phenotypic variance is minimized and covariance between reaction norm slope and phenotype is zero.

**Appendix S3** Equilibrium mean traits 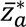 and 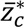 and equilibrium *u*_0_ when *P_ab_* = *P_bc_* = 0

**Appendix S4** Further mathematical analyses and comparisons with simulation results.

**Appendix S5** R code for simulations.

**Figure legends**

